# Optimization of Ethanol-Assisted Aqueous Oil Extraction from a *Cicadidae Sp*

**DOI:** 10.1101/2021.01.24.427958

**Authors:** Farzaneh Mahmoudi-Kordi, Mohammad Balvardi, Hamid-Reza Akhavan

## Abstract

Edible insects have been considering as a rich resource of high-quality protein and lipid content and at the same time a low-cost nutritious resource acquiring the least expenditure during farming, breeding, rearing and harvesting. On the other hand, organic solvent consumptions in industrial areas need to be limited; being flammable by environmental hazards. That is why seeking for alternative non-toxic solvents is highly substantial. In this study ethanol-aqueous extraction method applied at three different levels to find the optimum yield of edible oil extracted of Homoptera Cicadidae Magicicada, a seasonal insect residing underground, which is consuming locally as a whole meal. These levels set using Design of expert software to variables as pH, ethanol-concentration and solvent/sample ratio and the maximum yield extracted respectively at 6, 50 and 5, by the yield of 19.5%, that is comparable to oil extracted by conventional hexane oil extraction method at 27%. Fatty acids profile of recent method and hexane extraction method analyzed using GC-MS method and the physicochemical properties of either alternative method investigated. The results were on the standard ranges. Fatty acid profile of HCM was for the most part consisted of oleic acid, following by palmitic and linoleic acid 66.5, 19.2 and 7.8 respectively in hexane extraction method and 66.3, 17.7 and 9.5 in ethanol aqueous extraction method. The Iodine Value of HCM oil was noticeably high which introduces a medicinal applicatory besides the rich amount of MUFAs and PUFAs essential oils that are well-known for their incredible health benefits.

## Introduction

The United Nation Food and Agriculture (FAO) reported it is estimated by the year of 2050, world population will reach 9.1 billion. Leading that, the global food requirements will increase to 70% and this majority need to achieve more sufficient food and feed security.(Zielińska, Baraniak et al. 2015)Recently, scientists declared comparing to other sources of meat and poultry, edible insects can present a harmless and low-impact novel food and feed, owing to be such a nutritious source of protein, fat, micronutrients, minerals and fiber. (Huis, Itterbeeck et al. 2013, Yates-Doerr 2014) (Huis, Itterbeeck et al. 2013, Yates-Doerr 2014) (Huis, Itterbeeck et al. 2013, Yates-Doerr 2014) (Huis, Itterbeeck et al. 2013, Yates-Doerr 2014) (Huis, Itterbeeck et al. 2013, Yates-Doerr 2014) (Huis, Itterbeeck et al. 2013, Yates-Doerr 2014) (Huis, Itterbeeck et al. 2013, Yates-Doerr 2014) (Huis, Itterbeeck et al. 2013, Yates-Doerr 2014) (Huis, Itterbeeck et al. 2013, Yates-Doerr 2014) (Huis, Itterbeeck et al. 2013, Yates-Doerr 2014) (Huis, Itterbeeck et al. 2013, Yates-Doerr 2014) (Huis, Itterbeeck et al. 2013, Yates-Doerr 2014) (Huis, Itterbeeck et al. 2013, Yates-Doerr 2014) (Huis, Itterbeeck et al. 2013, Yates-Doerr 2014) (Huis, Itterbeeck et al. 2013, Yates-Doerr 2014) (Huis, Itterbeeck et al. 2013, Yates-Doerr 2014) (Huis, Itterbeeck et al. 2013, Yates-Doerr 2014) (Huis, Itterbeeck et al. 2013, Yates-Doerr 2014) (Godfray and Garnett 2014) From the environmental perspective, insect farms and breeding produce exponentially less greenhouse gasses, a practical approach to reverse the climate change cycle and global warming crisis, in addition to their minimal require of feed per unit and the rapid growth period, in comparison to livestock breeding.(van Huis and Oonincx 2017)

Approximately 1900 insect species for instance Mealworm, Grasshopper and Crickets are being traditionally appealing for human as a food resource, in African continent, Asia and Latin America.(Raheem, Carrascosa et al. 2019). In South Eastern region of Iran, non-periodical Homoptera Cicadidae *Magicicada* (HCM) is a familiar insect for locals using it as a meal after frying. This edible insect usually appears at surprising population amounts in springs. The nymphs reside underground in the enormous numbers together, sucking plant roots fluids to get fed. There are few studies focused on this spices of insect; However, past investigation revealed that it is a considerable resource of N, protein and lipids.(Brown 1973)

Various studies have considered edible insects as a substantial resource of protein and attended to technical functional properties, and extraction methods;(Chatsuwan 2018, Zielinska 2018)while less literatures focused on oil and fatty acids (FA) of which. Edible insects can be contributed as a different nutritional source of supplying and storing energy offering necessary fatty acids for human.(Ramos-Elorduy 2008) (Bukkens and Paoletti 2005) (Womeni, Linder et al. 2009, Xiaoming, Ying et al. 2010). This oil is usually rich of polyunsaturated fatty acids and the essential linoleic and α-linolenic FA that both of them are vital for a healthy diet, specially for children; Therefore, they can specially play a role at improving normal diet of people in developing countries with limited access to fish oil and sea foods.

In previous study by Li◻Feng. Yang et al on total lipid content and polyunsaturated fatty acid (PUFA) composition of six edible insect species (Mole Cricket, Ground Cricket, Spur◻Throated Grasshopper, Giant Water Bug, True Water Beetle and Water Scavenger Beetle), the predominant FA in all insects was oleic acid (18:1) and major SFA was palmitic acid (16:0), and also the main PUFA in most insects was linoleic acid (18:2n◻6). Furthermore, the ratio of n◻6/n◻3 was in range of 0.3 in spur◻throated grasshopper to 31 in Mole Cricket. In the another study, the main MUFA of many edible insects measured by E. Zielińska et al including adult Cricket (Gryllodes sigillatus), larvae of Mealworm (Tenebrio molitor), and adult Locust (Schistocerca gregaria), was palmitoleic acid (C16:1) and oleic acid (C18:1n9), and the uppermost content of oleic acid was specified in T. molitor at 40.86%. A. Adam Mariod et al also studied on edible oil extracted of Sorghum Bug (Agonoscelis pubescens) and the oil was 57.3% of dry bases, while the main fatty acids determined palmitic, stearic, oleic, and linoleic acids. Therefore, they showed a lower amount of saturated fatty acids and better content of overall unsaturated fatty acids.(Mariod 2020)

It is recommended to consider dietary fats as 20% - 35% of energy intake, with focusing on n-3 polyunsaturated fatty acids, and limiting saturated and trans fats(Vannice, Rasmussen et al. 2014). Fat consisted the main particle of total energy value, in whole components of edible insects. (Kouřimská and Adámková 2016) 7-77 percent of insect nutritional ingredient is consisted of fat, and this amount and the fatty acid compositions is related to their life stage as the oil concentration is higher in larvae, rather than adults, and also it is affected by their diet and the plants on which they feed.(Ramos-Elorduy, Moreno et al. 1997) (Fontaneto, Tommaseo-Ponzetta et al. 2011)

Demands on novel food industrial technology is rising these days; conversely, seeking a secure and efficient method to employ for edible oils extraction is a vital issue to remove technical barriers. Leading to the importance of this priority, alternative methods of extraction have been investigated during the recent decades. Many novel methods of oil extractions are suggested at laboratory scale as supercritical fluid extraction (SFE), pressurized liquid extraction (PLE), microwave assisted extraction (MAE), aqueous enzymatic extraction (AEA) and green solvents, etc. All the same, traditional methods like as expeller pressing or mechanical pressing and organic solvent are still more frequently used. (Kate, Singh et al. 2016)

n-Hexane, the nonpolar solvent prevailed of petrochemical sources, is the common organic solvent for oil extraction commercially; Owing to its feasibility in the industry, as low evaporation point, high consistency and stability, little corrosion and low residual oil content.(Toda, Sawada et al. 2016) And the result is a high quality oil, that the levels of free fatty acids (FFA) and anti-nutritional factors such as aflatoxins, gossypol and chlorogenic acids of that are less than other methods.(Hron Sr, Koltun et al. 1982)However, drawbacks as toxicity, flammability, time consuming process, labor-intensive must be considered. Moreover, it is such a hazardous material for health and ecosystems, and its volatile organic compounds, in case of emission to environment, can be contaminating to water and soil, and contributing to the troposphere ozone formation. (Kate, Singh et al. 2016)

In this context, alternative green and nontoxic solvents, including short chain alcohols, especially ethanol and isopropanol methods have been applied during solid-liquid extraction of lipophilic compounds.(Li, Fine et al. 2014, Martins and Peluzio 2015)

Combination of aqueous-ethanol, provides a polar protic solvent, that its ability of breaking up the emulsions, and high spreadability along the aqueous solvents, introduces an alternative method of oil extraction. While at the same time, it is renewable sources, worthy operating safety and low toxicity, that charges less than the other organic solvents. (Toda, Sawada et al. 2016) Tatiane Akemi Toda et al. studied on kinetics of soybean oil extraction using absolute ethanol and aqueous ethanol (5.98mass% of water) as solvent in different temperatures (40, 50 and 60 °C). The result showed that the water level in the solvent decreased the soybean oil extraction while increasing values of temperature favored the oil extraction and free fatty acids.(Toda, Sawada et al. 2016)

As it is mentioned, there is lack of studies focusing on insect’s oil as a novel sources of energy and also on nutritional properties of HCM, the edible insect; Therefore, the primary aim on this study was the modality and functionalities of C.H oil, aiming to eliminate consequences of conventional organic solvents. SO the green ethanol aqueous extraction (EAE), established as the extraction method. Thereupon, the secondary aim was finding the optimum yield of HCM oil, extracted by this method, that evaluated under different levels of variables as pH, ethanol concentration and solvent/sample ratio. Simultaneously, parameters as approximate properties analyses, oil color, physicochemical characterizations, and fatty acids profile of the of H.C oil, extracted by recent novel method, analyzed and compared to those of the conventional hexane-extraction method and the recommended standard amounts.

## Materials and Methods

### Materials

Cicadas are winged, plant-feeding insects of the order Homoptera, that are active through the evening, and usually are not considered as pests. Typically, greatest amount of cicadas in a *Magicicada* population appear in synchrony in surprising bulks of populations, that generally appear in May, June and July, and stay alive for a few weeks. Cicadas have multiple-year life cycles and most of them are non-periodical which means the adults are existing every year, whereas the others are partial periodicity.(Marlatt 1907, Marshall 2001)

500 g wild adult Homoptera Cicadidae *Magicicada* (photo at figure 1) were purchased from local store in Kerman-Iran in July 2020 and transformed to laboratory. All chemical materials and solvents were on analytical grade of Merck Chemical Company to be used in chemical analyses,(O’Neil 2013) and analyses were conducted in triplicate (n = 3).

**FIGURE 1.**
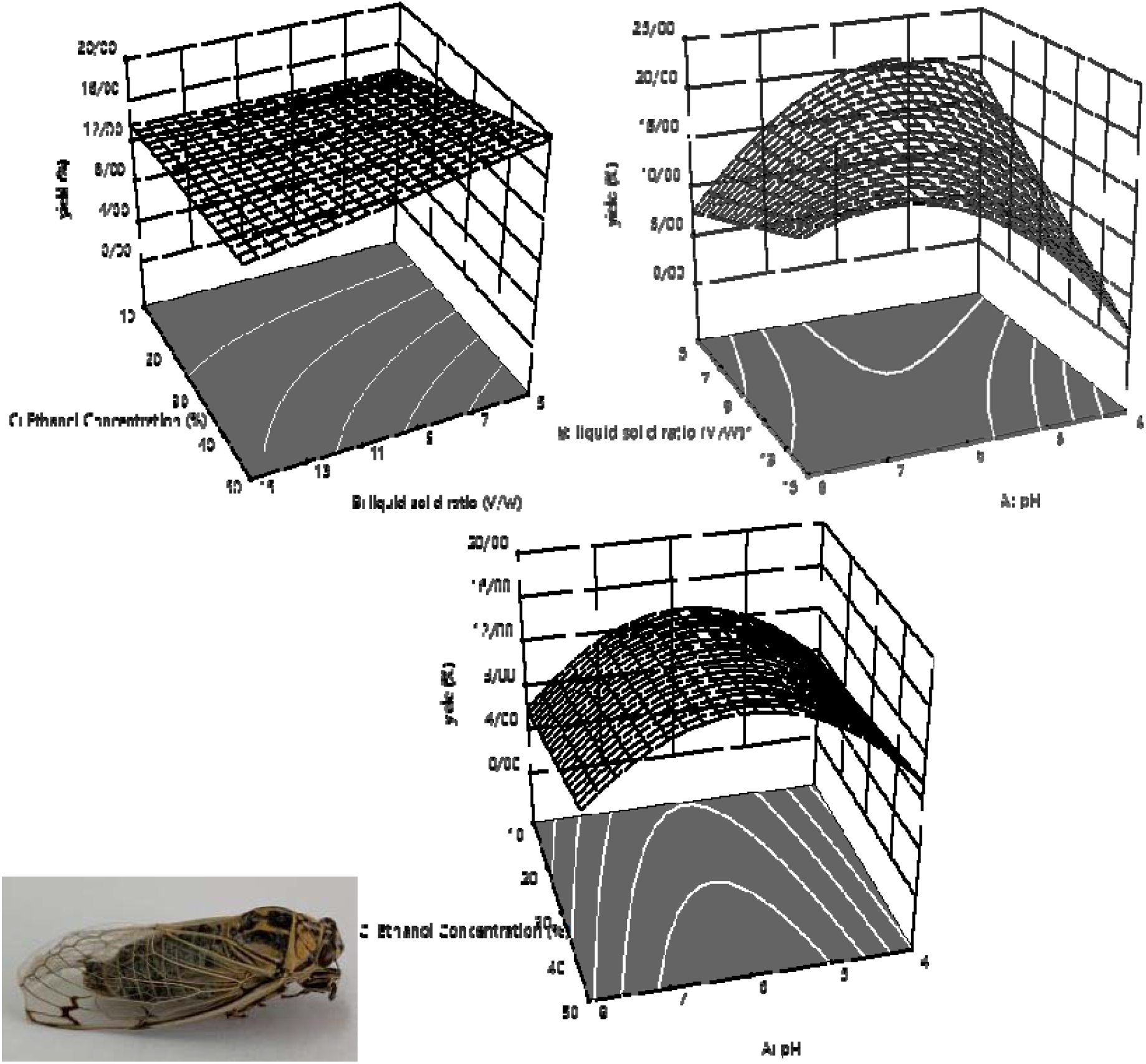
Frozen Homoptera:Cicadidae:Magicicada, The adult insect.

### Methods

#### Aqueous Ethanol Extraction Procedure

HCM live insects starved a day, for the reason that their gastrointestinal tract get empty, then froze and killed by liquid Nitrogen and subsequently freeze-dried (China Laboratory Freeze Dryer, MC-FD10B), according to S. Ghosh et al. (Ghosh, Chuttong et al.). Powder-lyophilized samples, sieved through a 30-mesh and packed into plastic vacuumed pockets and stored in −20.0°C freezer to prevent the enzymatic degradations and physicochemical changes, for further experiments. Three levels of solvent/sample ratio, ethanol concentration and pH provided, according to the data of Design of Expert software showing at table 1. Leading that, determined volume of ethanol (96% excipient) and deionized water mixed on decided ratio, and each portion of solvent added to its samples in labeled Falcon tubes, at triple repetitions. Afterwards, samples shook using laboratory shaker (KS,260 Basic) at 300 rpm and 15 min, and pH adjusted using digital desktop pH meter (Shimaz, PTR79) using NaOH 1N and HCL 0.1N. Simultaneously, samples shook again and incubated in hot water bath (IKA, HB digital) at 50°C for 30 min. To finalize oil extraction, and separate the oily phases, samples centrifuged (Sigma,3-16KL) at −4°C, 4500×g for 20 min. The oily phase separated completely on the top of the other particles after centrifuging, and then transferred to plates, using an automatic micropipette (New Cayon, 10-100μl). The weight of clear oil reported to calculate the oil extraction yield, according to the following equation and extracted oil samples were froze at −20 °C for further analysis.(Tzompa-Sosa, Yi et al. 2014)

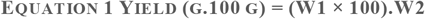

**TABLE 1.**
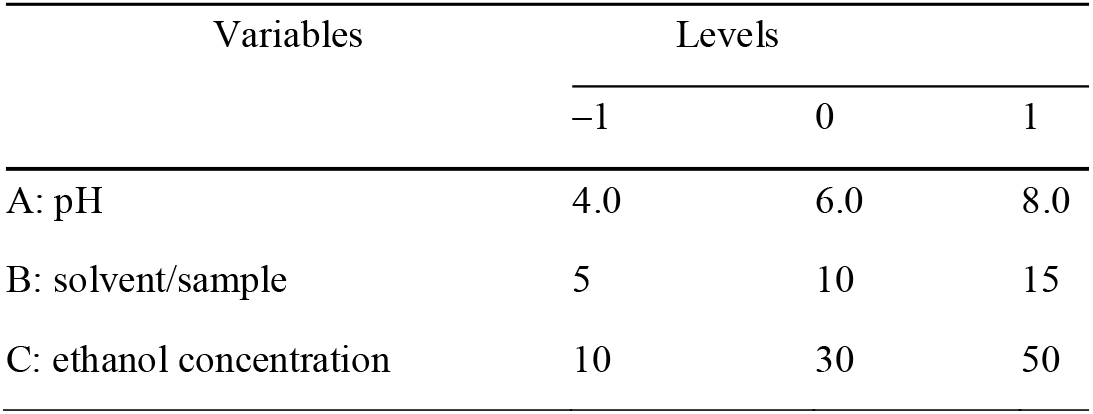
CODED AND ACTUAL VARIABLE LEVELS FROM THE BOX–BEHNKEN DESIGN USED IN THIS STUDY FOR THE AQUEOUS-ETHANOL EXTRACTION OF OIL

| Variables | Levels |  |  |
| --- | --- | --- | --- |
|  | −1 | 0 | 1 |
| A: pH | 4.0 | 6.0 | 8.0 |
| B: solvent/sample | 5 | 10 | 15 |
| C: ethanol concentration | 10 | 30 | 50 |

Where, W1 is the weight of the separated clear oil from the supernatant, and W2 is the weight of the whole insect powder used for extraction. figure 2 illustrates the schematic process of extracting oil through the recent method.

**FIGURE 2.**
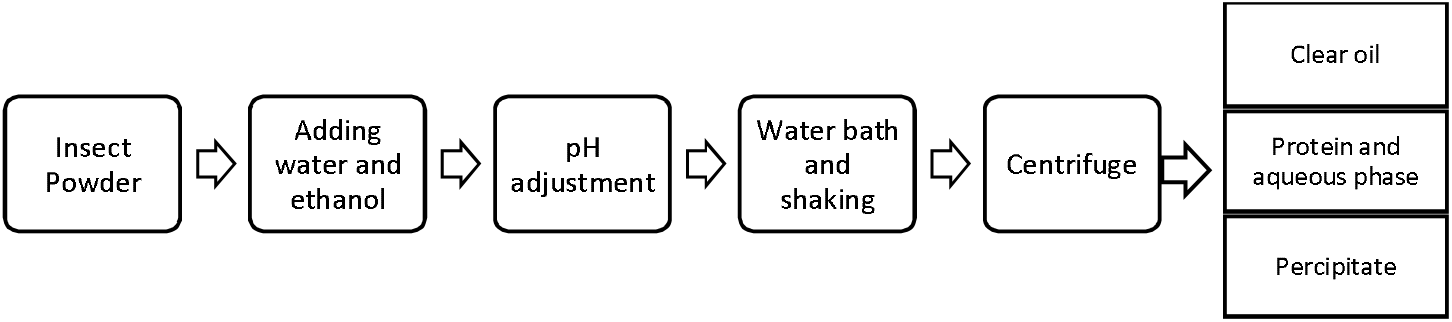
Schematic representation of aqueous-ethanol extraction ofHCMOil, using three levels of Solvent/Sample ratio (5, 10, 15 ml.gr), ethanol concentration (10, 20, 30 %) and pH (4, 6, 8) and three phase of the final result after centrifuging.

### Experimental Design for Response Surface Methodology

Box–Behnken design (BBD) (Zhang, Zhang et al. 2009) used to obtain the highest oil recovery, at three levels of independent variables as pH (A), solvent/sample ratio (ml/g, B), ethanol concentration (ml, C), which respectively set at 4.0, 6.0, 8.0 and 5, 10, 15 and 10, 30, 50 levels, as it is demonstrated on table 1, and the mean values of oil recovery (%) were considered as the response. Regression analysis of the data, was performed according to the experiential second order polynomial model, as stated at the following equation (Balvardi, Rezaei et al. 2015) (Zhang, Zhang et al. 2009)

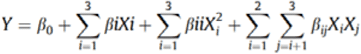

Where *Y* demonstrates oil recovery and β_*0*_, β_*i*_, β_*ii*_ and β_*ij*_ are respectively the regression coefficients in the intercept, linear, quadratic, and interaction terms besides *X_i_* and *X_j_* that are independent variables. Design of Expert (DOE) software version 11 (Stat-Ease Inc., Minneapolis, MN) applied to discover coefficients of the quadratic polynomial model.

### Hexane Extraction Method

Aiming to extract HCM oil by Sohxlet method(AOCS 1998), as the control method for determination the recoveries obtained by EAE method, the insect powder provided as it stated before. Then, it sieved through a 30 μm laboratory mesh, and 50 g weighted sample by a laboratory balance (Sartorius, GE412) wrapped through a filter paper, and inserted into Soxhlet extractor equipment. 250 ml n-hexane (1:5 gr/ml of sample/solvent ratio) added into an Erlenmeyer boiling flask (250 ml) connected to the condenser, and heated by a heating mantle (Wisd, WiseTherm) at 50 °C for 6 h, to complete oil extraction procedure. Ultimately, solvent removal procedure applied using rotary evaporator (IKA, RV 10 digital V) to pursue the pure oil until no solvent was seen, and oil samples were froze at −20 °C for further analysis.(Tzompa-Sosa, Yi et al. 2014)

#### Proximate Analyses

Chemical compositions were analyzed, using approved methods of standard official methods of analysis of AOAC international(international 2006). Physicochemical properties as crude protein determined using the Kjeldahl method, (AOCS 1998) and a nitrogen-to-protein conversion factor of 5.95 was considered (Int 2007) for accounting total proteins. The fat percentage was calculated by extraction in a Soxhlet using hexane as the solvent (AOCS 1998). Ash content measured by drying the samples in a muffle oven (550 °C for 5 h) (Int 2007), and the moisture analyzed by drying the samples into an oven (105 °C for 2 h) (AOCS 1998). The carbohydrate content was estimated by difference, according the following equation: 100 - (weight in grams [protein + fat + moisture + ash] in 100 g of edible insects).(Kekeunou, Simeu-Noutchom et al. 2020)

### Physicochemical Properties

Parameters as refractive index (RI), acid value (AV), Iodine Value (IV), saponification value (SV), unsaponifiable matter (UM), peroxide value (PV), para-anisidine value (*p*-AV), and TOTOX value of EAE method, measured and compared to those of oil HE method. RI, IV, SV and UM, were determined using the methods of 921.08, 920.158, 920.160 and 933.08 respectively, according to Official Methods of Analysis of AOAC International (AOAC 2000). The AV, PV and *p*AV were determined by methods of Ca 5a-40, Cd 8b-90 and Cd 18-90, respectively, from the AOCS Official Methods of Analysis (Bindhu, Reddy et al. 2012)and also The TOTOX value was determined according to the equation of 2 × PV + *p*-AV. (Poulli, Mousdis et al. 2009) Energy content calculated according to the following organic matter (1 g of carbohydrate yields 4 kcal, 1 g of lipid = 9 kcal, 1 g of proteins = 4 kcal in the bomb calorimetric)

### Color Quality

One of the most important elements in industrial refining process and also in marketing is the color of edible oils.(Fengxia, Dishun et al. 2001) To this aim, a combination of digital camera, computer, and graphics software set up to measure and determine the color of HCM oil, based on the L*a*b* format according to (AOCS Cc 13c-50)(Sandulachi and Tatarov 2014). Instrument calibration was set by the zero calibration box and the white calibration plate, used for total color difference calculation.

#### Fatty Acid Analysis

Gas chromatography is considered as a delicate, precis, accurate quantitative and qualitied method to analysis complex fatty acid profile of oils(Van Ruth 2001). Fatty acids esters (FAEE) privatized by the method of Golmakani and to this purpose, 30 mg of both samples of oil were treated with 3.0 mL of ethanol-acetyl chloride to the ratio of 95:5 v.v and mixed well under a nitrogen atmosphere.(Balvardi, Rezaei et al. 2015) The temperature increased to 85 C for 1 h and after cooling to the room temperature, 1.0 ml double distilled water added to a clean vial and shook for 1 min. Then 3.0 mL of n-hexane added to provided FAEE solution and transferred to a vial and injected to GC-MS for further examinations. To analyze FFA a split.split-less injector coupled to a QP-2010 plus single quadruple mass spectrometer and for separating FFA a 007-CW Carbowax fused silica capillary column (12 m × 0.1 mm I.D. × 0.1 μm film thickness) from Quadrex (Woodbridge, CT) operated. In the injector, interface and ionization chamber, the temperature were maintained at 220, 240 and 230 °C, respectively. The temperature of oven set up to start at 100 °C, heated to 160 °C at a rate of 20 °C.min and continuously reached to 220 °C at a rate of 15 °C.min. At this point, it held for 8 min and Helium at a flow rate of 0.8 ml.min was considered as the carrier gas. A portion of 0.5 μL of the sample was ultimately injected into the GC–MS while the injector programmed in the split mode (split ratio 1.10) and compounds were primarily recognized, by mass spectrometry in the SCAN mode, by a mass interval ranging of 40 to 400 m.z. FAEE, were finally categorized by comparing their retention times to those of the standards.

### Statistical Analysis

All experiments completed by the BBD and were statistically analyzed using Design of Expert software. All the further quality analysis experiments, applied to the extracted oils, were conducted in triplicate and statistical analysis was performed using student’s t test from SPSS at a p < 0.05 significance level.

## Result and Analysis

### Optimization of Oil Recovery

The optimized conditioned (coded) and observed (actual) values of EAE via different variable combinations, according to the Box–Behnken design, at 17 run is presented at table 3. The effect of the pH on the oil recovery in oil aqueous extraction previously investigated by Y.B. Che Man et al, on aqueous enzymatic extraction of coconut oil, that stated there were significant increases at oil yield by increasing pH from 4 to pH 7; However, it decreased after, by reaching to pH 8 again(Man, Asbi et al. 1996). At the series of recent experiments, this trend observed by increasing the yield up to pH 6, and decreasing reaching to pH 8. So that, three levels of pH set at 4, 6 and 8. The influence of different ratios of the solvent/sample on the oil extraction yield, was studied at 5:1, 10:1 and 15:1 levels at the previous study by Q.-A. Zhang et al, on autoclaved ultrasound-assisted extraction oil of almond powder. The results showed a considerable increase at the amount of extracted oil, by increasing this ratio. Also it is in agreement with the mass transfer principle, that states the driving force during the mass transfer is the concentration gradient, among the solid and the volume of the liquid, while it is greater when solvent/sample ratio increases(Zhang, Zhang et al. 2009). The three levels of variable concentrations of ethanol using for extraction were determined according to the results of a series of experiments, from absolute deionized water to 50% ethanol concentration, that it was considered as the maximum level, to find out the highest economical yield of extraction by using at least ethanol portion, and according to the results, 10%, 30% and 50% set as the variables. At pH 6 and solvent/solid ratio 5:1, oil extraction yield increased substantially, by increasing the ethanol concentration at solvent, as the oil recovery reached to the maximum point at the yield of 19.5% by 50% ethanol concentration. This behavior previously reported by Toda et al, studying on soybean oil extraction, using ethanol as solvent, that stated decreasing aqueous phase in ethanol, increases the yield of oil during extraction(Toda, Sawada et al. 2016). As well as it was also previously mentioned at another study, on rice bran oil using ethanol as solvent. (da Costa Rodrigues, Oliveira et al. 2010)

Table 2 shows the results of analysis of variance for experiment design of the Box–Behnken, and corresponding response data of the oil extracted by EAE method. The coefficient of predicted R2 and adjusted R2 of ANOVA statistics results, were determined at 0.9896 and 0.9948 respectively, that suggests the quadratic model. Furthermore, the model F-value, 342.86, inferred that it was significant; Beside that, results of the error analysis, indicated that the lack of fit was insignificant (p > 0.05). The final expression in terms of coded factors is given as following equation.

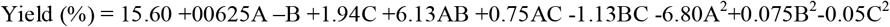

**TABLE 2.** ANALYSIS OF VARIANCE (ANOVA) FOR THE CODED FACTORS OF OIL RECOVERY BY AQUEOUS-ETHANOL ALMOND IN THE CURRENT STUDY

| Source | Sum of Squares | df | Mean Square | F-value | p-value |
| --- | --- | --- | --- | --- | --- |
| Model | 391.23 | 9 | 43.47 | 342.86 | 0.0001> significant |
| A-pH | 0.03 | 1 | 0.0313 | 0.25 | 0.6348 |
| B-liquid solid ratio | 8.00 | 1 | 8.00 | 63.10 | 0.0001> |
| C-Ethanol Concentration | 30.03 | 1 | 30.03 | 236.87 | 0.0001> |
| AB | 150.06 | 1 | 150.06 | 1183.59 | 0.0001> |
| AC | 2.25 | 1 | 2.25 | 17.75 | 0.0040 |
| BC | 5.06 | 1 | 5.06 | 39.93 | 0.0004 |
| A<sup>2</sup> | 194.69 | 1 | 194.69 | 1535.62 | 0.0001> |
| B<sup>2</sup> | 0.02 | 1 | 0.03 | 0.19 | 0.6786 |
| C<sup>2</sup> | 0.01 | 1 | 0.02 | 0.08 | 0.7816 |
| Residual | 0.89 | 7 | 0.13 |  |  |
| Lack of Fit | 0.19 | 3 | 0.07 | 0.36 | 0.7880 not significant |
| Pure Error | 0.70 | 4 | 0.18 |  |  |
| Cor Total | 392.12 | 16 |  |  |  |

**TABLE 3.**
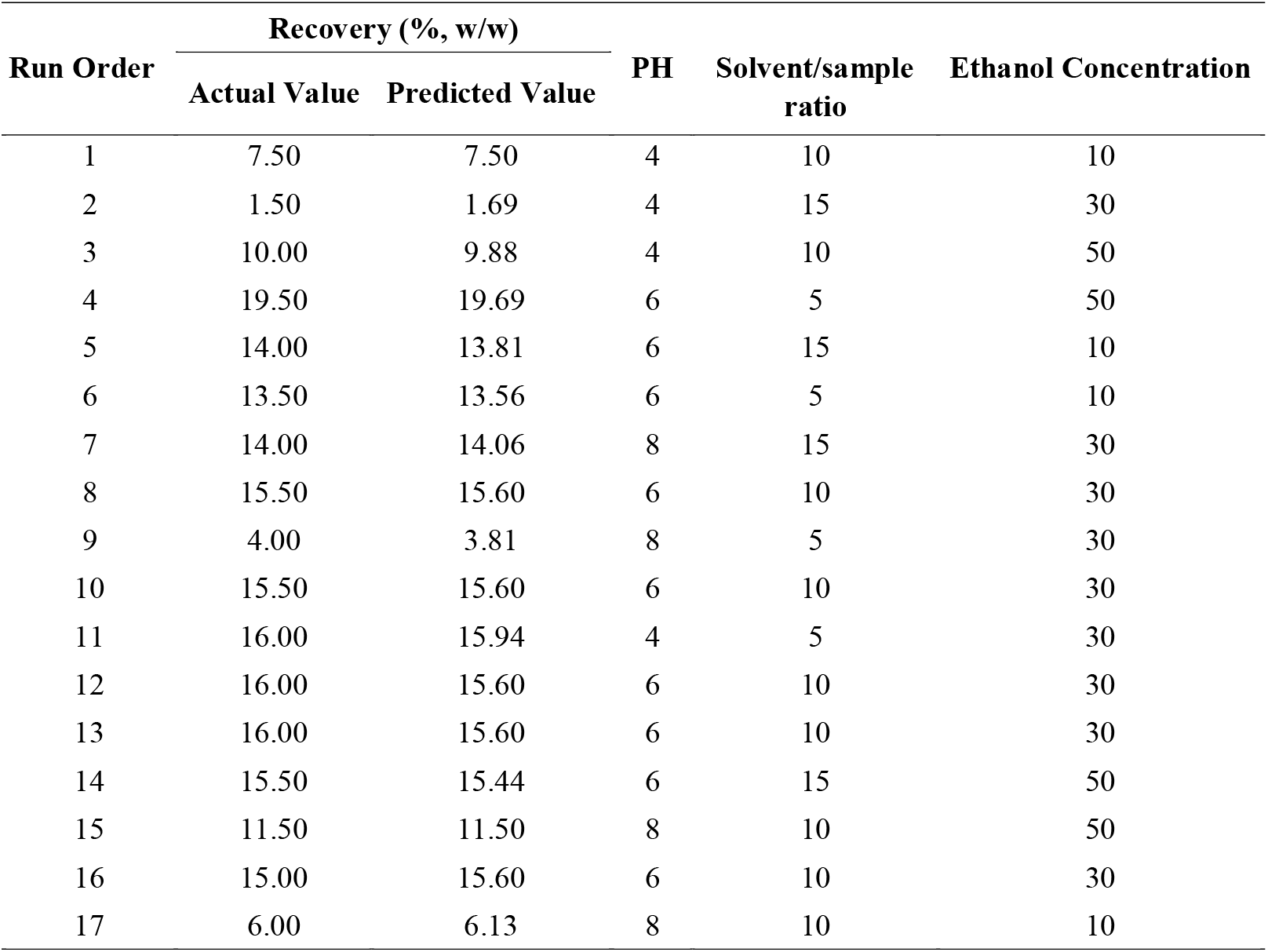
Obtained o]il recoveries at the various conditions applied in the current study for the aqueous-ethanol extraction of oil from Iranian wild almond compared with the predicted values based on the equation suggested by Box–Behnken design

| Run Order | Recovery (% w/w) |  | PH | Solvent/sample ratio | Ethanol Concentration |
| --- | --- | --- | --- | --- | --- |
|  | Actual Value | Predicted Value |  |  |  |
| 1 | 7.50 | 7.50 | 4 | 10 | 10 |
| 2 | 1.50 | 1.69 | 4 | 15 | 30 |
| 3 | 10.00 | 9.88 | 4 | 10 | 50 |
| 4 | 19.50 | 19.69 | 6 | 5 | 50 |
| 5 | 14.00 | 13.81 | 6 | 15 | 10 |
| 6 | 13.50 | 13.56 | 6 | 5 | 10 |
| 7 | 14.00 | 14.06 | 8 | 15 | 30 |
| 8 | 15.50 | 15.60 | 6 | 10 | 30 |
| 9 | 4.00 | 3.81 | 8 | 5 | 30 |
| 10 | 15.50 | 15.60 | 6 | 10 | 30 |
| 11 | 16.00 | 15.94 | 4 | 5 | 30 |
| 12 | 16.00 | 15.60 | 6 | 10 | 30 |
| 13 | 16.00 | 15.60 | 6 | 10 | 30 |
| 14 | 15.50 | 15.44 | 6 | 15 | 50 |
| 15 | 11.50 | 11.50 | 8 | 10 | 50 |
| 16 | 15.00 | 15.60 | 6 | 10 | 30 |
| 17 | 6.00 | 6.13 | 8 | 10 | 10 |

Where A, B and C respectively indicate pH, solvent/sample ratio and ethanol concentration. As it is obvious, A, C, AB, and AC positively affect the response, while the rest of the parameters have a negative influence.

According to mentioned equation, optimal levels of the variables for the oil extraction of HCM constructed at three-dimensional surface plots as it is shown at figure 2.

As graph (a) demonstrates, the interaction of pH and solvent/sample ratio positively effects the yield. By increasing pH and solvent/sample ratio, respectively from 4 to 8 and 5 to 15, the yield increases considerably; However this trend for pH is also increasing by reaching to 5.87 (the pick of graph), but it decreases slowly by pH 8. Besides, solvent/sample ratio keeps the increasing trend moderately from 5% to 15% all the way long. Graph (b) shows ethanol concentration and sample/solvent ratio across the yield and their interactions. There was a considerable increase by growing the ethanol concentration amount and also an adequate decrease in the yield by rising sample/solvent ratio. Likewise, the interaction of these two variables was also following a decreasing trend too. The last graph, (b) shows the yield through pH and ethanol concentration. By increasing pH, firstly the yield increases moderately to around pH 6.0, and then drops reaching to pH 8.0. Although, the trend of ethanol concentration is substantial rising from 10 to 50 % ethanol concentration, and their interception follows a moderate increasing trend.

The optimal conditions anticipated by BBD determined at pH 6.0, ethanol concentration 50% and solvent/sample 5:1 which leads to an oil recovery of 19.688%. To compare the predicted result with the actual value, the experiments checked again, according to inferred optimal condition. The mean value of 19.50 ±0.33% (n = 3), observed of the real experiments, that validated the RSM model (p > 0.05).

As it mentioned, the maximum yield of ethanol aqueous extraction determined at 19.50 ±0.33% experimentally, and also due to the results of table 4 the oil extracted by Soxhlet method was 27 ±0.18%; Therefore, there was a significant difference between these two methods. The extracted yield (%) was calculated based on total lipid content of EAE/HE at 72.22 ± 0.18 %, which is comparable to previous findings of D.A. Tzompa-Sosa et al on aqueous versus organic solvent-based extraction oil of some insect spices, that this ratio reported at 61.41, 51.40 and 40.78 respectively for T. molitor, A. diaperinus and B. dubia(Tzompa-Sosa, Yi et al. 2014). However, this ratio in the recent study was higher, according to positive effect of adding ethanol to aqueous solvent, in comparison to using water simply as solvent.

**TABLE 4.**
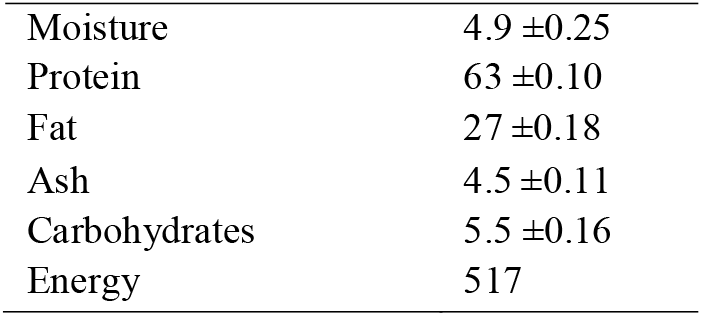
CHEMICAL COMPOSITION OF HOMOPTERA CICADIDAE (G.100G DRY MATTER). VALUES ARE MEANS ± SD OF TRIPLICATE DETERMINATIONS.

**Figure 2** Dimensional surface plots of the combined effects of pH and solvent/sample ratio (a) and pH and ethanol concentration (b) and ethanol concentration and solvent/sample ratio (c) (gr/ml) each (shaking rate: 300 rpm)

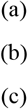

#### Approximate Properties

The approximate properties of HCM powder in 100 g dry matter, and also the moisture content are demonstrated on table 4. These figures show that the greatest portion of nutrients is for protein content with 63±0.10% of total dry matter. This high protein content is in agreement to the average crude protein content of insects, varying from the range of 40 to 75% dry weight. (Verkerk, Tramper et al. 2007, Rumpold, Schlüter et al. 2013)This large protein scale source can provide a substantial contribution to prevent protein deficiency, especially in developing countries, and meet the needs of infant growing and well-being to fight against protein malnutrition. HCM is also an important source of fat by 27 ±0.18 % oil, which is a considerable higher than the other edible insect species as *Kraussaria angulifera* and *Acanthacris ruficornis*that that respectively are consisted of 11.71 and 9.00% oil, and comparable to species as *Hemiptera* (true bugs) and *Lepidoptera* (butterflies, months) by respectively 30.26 and 27.66% fat. The ash and Carbohydrate content also measured at 4.5 ±0.11 and 5.5 ±0.16, while the moisture content was determined at 4.9%. The result shows that 100 g of HCM powder provides 517 kcal energy that is comparable to same amount of fresh meat saving pork meat which may be more as the result of extreme fat content.(Sirimungkararat, Saksirirat et al. 2010)

#### Physiochemical Characteristics of the Oil

The commence of industrial aspiration factors of edible oils and fats, is their functional properties, since it is related to the physicochemical and structural properties conferring to the final product. (Belton 2000) To monitor edible oil specifications, in case of insuring the consumption qualities, some paramount physiochemical characteristics studied and the results are presented on the table 5.

**TABLE 5.** Physicochemical characterization ofHCMOil Results presents the mean ± SEM of three estimations, Appearance is based on physical states at room temperature (25°C), RI refractive index, FFA free fatty acids value (mg KOH.g oil),

| Quality Parameter | Hexane Extraction | Ethanol-Aqueous | FAO.WHO Standards |
| --- | --- | --- | --- |
| Yield (%) | 27 ±0.18 | 19.688% |  |
| Color | Deep yellow | Yellow |  |
| Appearance | Liquid | Liquid |  |
| RI | 1.4630 | 1.4632 | 1.467-1.470 |
| FFA | 2.006 ±0.13 | 1.494 ±0.55 | 4 |
| IV | 89.96 ±0.22 | 97.15 ±0.51 | 84-98 |
| SV | 186.32 ±0.25 | 193.33 ±0.72 | 187-196 |
| USM | 8.11 ±0.80 | 8.74 ±0.02 | ≤30 |
| PV | 0.21 ±0.65 | 0.28 ±0.18 | 10 |
| p-AV | 5.61 ±0.2 | 6.63 ±0.11 | 20 |
| TOTOX | 6.03 ±0.50 | 7.19 ±0.14 | 26 |

Free fatty acid portion at the method of EAE determined at 1.494 ± 0.55, that was considerably less than that of HE method at 2.006 ± 0.13. Subsequently, the amount of free fatty acid had lessened during the recent method that is in the agreement to the previous study by D.A. Tzompa-Sosa et al, that reported less FFA in aqueous extraction method in several insect spices. (Tzompa-Sosa, Yi et al. 2014) In another study, Hron Sr et al. also implied that using ethanol as the solvent, reduces the amount of total FFA during the extraction process rather than Hexane as solvent.(Hron Sr, Koltun et al. 1982). This can be related to the fact that FFA form soaps in the presence of sodium or calcium salts, that are common micronutrient of insect structures, that leads to increasing FFA solubility in water. So, they would have been washed out by aqueous phase, and separated from the oil on the supernatant in EAE. (Tzompa-Sosa, Yi et al. 2014) This reduction in FFA proves that it is significantly affected by the solvent type, and EAE method provides an oil with higher oxidation stability, that eliminates further refining steps, and as result, reduces extra energy consumptions and further costs. (Toda, Sawada et al. 2016)

Saponification value in EAE method was measured at 193.333 ± 0.55 and in HE at 186.316 ± 0.25. So it is implied that the molecular weight and the concentration of fatty acids extracted by aqueous ethanol solvent was way more than those of hexane extraction. (Bart, Palmeri et al. 2010) Whereas there was no significant difference between unsaponification matter that respectively determined in EAE and HE methods at 8.74 ± 0.02 and 8.11 ± 0.8.

Iodine Value (IV) in EAE method was determined at 97.15 ± 0.51 (g I/100 g) while in the other method it was 89.96 ± 0.22 (g I/100 g) which is comparable to this amount in black soldier fly larvae at 90.1 (g I/100 g) (Su, Nguyen et al. 2019) and also in another study of mealworm oil at 96 (g I/100 g) (Mariod 2020). It can be contributed that unsaturated fatty acids which relating to melting point and oxidative stability, were higher in alcoholic solvent rather than hexane. (Dijkstra 2016) High Iodine Value can also predict a high medicinal value, as it is an index of the degree of unsaturation of the oil, according to Ekpo et al that reported APW larval oil has a great amount of IV (108 ± 0.15 to 140 ± 0.51), that leads to pharmaceutical potential(Ekpo, Onigbinde et al. 2009)

Turning to oxidation indicators, such as the peroxide value (PV), that in HE and EAE methods determined 0.21 ± 0.65 meq O_2_.kg oil and 0.28 ± 0.18 meq O_2_.kg oil, respectively. There is no significant difference at using either of these solvents, in the amount of initial oxidation products, as in previous study by Belssem Jedidi et al, on Tecoma stans seed oil, the amount of peroxide value, while using either hexane or ethanol solvents during extraction, was roughly similar.(Jedidi, Mokbli et al. 2020). P-Anisidine value (P-AV) and TOTOX respectively were 5.61 ± 0.2 and 6.03 ±0.50 in HE and 6.63 ± 0.11 and 7.19 ± 0.14 in EAE. As the result shows, P-AV is noticeably higher in EAE which reveals that the amount of secondary oxidation products are considerably higher at EAE method, as a consequence of the oxidation products, which are more miscible in polar solvents as ethanol rather than un polar hexane and therefore they extracted in greater scales by the ethanol aqueous solvent.(Small 1953)

Physical characterizations as color and appearance in HE and EAE method respectively were deep yellow and yellow, and liquid physical stated in both of extraction methods at the room temperature (25 °C). Refractive index (RI) in the HE methods was 1.4630 and in the other method measured at 1.4632 which indicates that there were no significant differences between these two samples.

### Fatty Acid Profile

Table 6 illustrates the summarized results from GC-MS, the type of fatty acids and the quantities of HCM oil extracted by two different methods of hexane extraction method HE and EAE. At the first glance, it is noticeable that the total amounts of fatty acid in both extraction methods is roughly equal with more or less differences.

**TABLE 6.** Fatty Acid Profile of *homoptera cicadidae magicicada*

| Fatty acids | H.C Hexane<br>extraction (%) | H.C Aqueous<br>extraction (%) |
| --- | --- | --- |
| Palmitic C16:0 | 19.2 | 17.7 |
| Palmitoleic C16:1 | 1.4 | 1.6 |
| Stearic C18:0 | 3.2 | 2.6 |
| Oleic C18:1 | 66.5 | 66.3 |
| Linoleic C18:2 | 7.8 | 9.5 |
| Linolenic C18:3 | 1.1 | 0.6 |
| Arachidic C20:0 | nd | 0.6 |
| Behenic C20 | 0.2 | 0.7 |
| Lignoceric C22 | 0.2 | 0.4 |
| SFA | 22.8 | 22.0 |
| USFA | 76.8 | 78.0 |
| MUFA | 67.9 | 67.9 |
| PUFA | 8.9 | 10.1 |
| PUFA/SFA ratio | 0.39 | 0.45 |
| MUFA/PUFA ratio | 7.62 | 6.72 |
| SFA/USFA | 0.29 | 0.28 |

The result shows that HCM oil is a C18 carbon-rich oils, with total C18-carbon fatty acid content of 78.6 and 79 percent at HE and EAE respectively. Among the C18-carbon fatty acid amounts, oleic acid (C18:1) was the dominant fatty acid, varying from 66.5% at HE to 66.3% at EAE. This large scale of oleic acid is comparable to fatty acid composition of olive oil, that mainly consists of oleic acid (ranging from70–80%)(Sales-Campos, Reis de Souza et al. 2013).

Simultaneously, in some species of edible insects in previous studies, it was stated that oleic acid determined the most dominant fatty acid. As in a study by Sihamala et al on black ants (*Polyrhachis vicina*) oleic acid was ranging from 52.1 to 63.0% (Oranut, Subhachai et al. 2010) and in another study by Sampat Ghosh et al on Asian honeybee species (*apis cerana and apis dorsatabee*), it was at 57.68% (Ghosh, Chuttong et al.) and also in the other study by Igwe et al on African termites, it measured at 52.45% of total FAs. (Igwe, Ujowundu et al. 2011).

Accordingly, it can be subsidized that to date, the uppermost content of oleic acid in different edible insect species, has been found at HCM oil, and it is a great source of this vital fatty acid. From a nutritional point of view, oleic acid plays a potential role at decreasing the risk of coronary heart disease(Michaelsen, Hoppe et al. 2009), and also it has positive effects on the autoimmune system, countering the inflammatory diseases (Sales-Campos, Reis de Souza et al. 2013) and reducing brain-related disorders, for instance Alzheimer’s disease and dementia (Aluko 2012) that must be considered in a healthy diet and HCM as an reach oleic acid sours, can meet this need.

The second principal fatty acid was palmitic acid (C16:0) which in HE and EAE method respectively measured at 19.2 and 17.7, that showed it was dominated in HE method and roughly near to this fatty acid percentage in black ants ranging from 16.5 to 20.8%. (Oranut, Subhachai et al. 2010) Among the other C16-carbon fatty acids, palmitoleic acid (C16:1) varied from 1.4% in HE to 1.6% in EAE.

Linoleic acid C18:2 extracted by EAE solvent was 9.5% and comparably higher than that of HE at 7.8%, and linolenic C18:3 amount was also 0.6 and 1.1% respectively. Therefore, this oil is rich in polyunsaturated fatty acids and essential fatty acids, which cannot be synthesized by human body that must be considered in diet because of their essential role on normal growth and dermal function specially in children.(Holman 1998, Michaelsen, Hoppe et al. 2009) However, people in developing countries with restricted access to sea foods, are experiencing possible deficiencies, and potential lack in intake of these two essential polyunsaturated fatty acids.(Cuvelier, Cabaraux et al., Michaelsen, Hoppe et al. 2009, Roos, Nurhasan et al. 2010, Huis, Itterbeeck et al. 2013). Therefore, HCM oil can conveniently be used as an alternative to attain a food supplement and nutritionally rich diet, and using ethanol aqueous as the solvent will increases the amount of these essential fatty acids.

According to these data, the lipid levels at HCM are comparable to fishery products, as the amount of its saturated fatty acids (22.8% and 22.0% in HE and EAE methods respectively) are similar to those of wild Atlantic Salmon (19.0%) and Rainbow Trout (24.4%), and also the edible insect Sorghum Bug (20.8%)(Blanchet, Lucas et al. 2005, Mariod 2020)while this portion at Beef cattle (44.0%) and chicken (34.7%)(Rule, Broughton et al. 2002) is excessively high, that especially in red meat has been associated to risk of numerous major chronic diseases as diabetes, coronary heart diseases and cancer.(Chakravarthy, Jayasimha et al. 2016, Wolk 2017)

Furthermore, HCM oil by around 78% USFA at both methods rich in unsaturated. As D.A. Tzompa Sosa reported, the insect oil studied in their laboratory, were rich in unsaturated FA (>70%) too (Tzompa-Sosa, Yi et al. 2014) that was the reason of the liquid appearance even at low temperatures. On the other hands, this could limit their application in food industry, as different food formulation requires flexible physicochemical properties. However, this issue can be tackled by changing physical properties using dry fractionation, and altering the chemical composition.(Wassell and Rajah 2014)

The other fatty acids as stearic, palmitoleic, behenic, and lignoceric acids respectively at HE method were 3.2, 1.4, 0.2 and 0.2%, and at EAE method 2.6, 1.6, 0.7 and 0.4 %.

While there was no Arachidic acid detected in HE method, the amount of 0.6% of that extracted by EAE. It can be extracted that wide spectrum of fatty acids extracted by EAE rather than HE method and this can be allocated as an advantageous aspect of recent method. In edible insects, the range of saturated to unsaturated fatty acids is from 0.4% to 0.65%(Mariod 2020). By increasing unsaturated fatty acid amounts this ratio decreases as in the recent study this ratio was 0.29 that is on the range of previous investigations.

Nevertheless, the previous study on cicada (meimuna opalifera walker) by Raksakantong et al. indicated that its fatty acid profile was completely different from those of homoptera cicadidae magicicada in recent study, and predominated fatty acid of that consisted of stearic acid, C20:4n6 and C20:3n6 respectively ranging 52.53, 33.03 and 10.77%.(Raksakantong, Meeso et al. 2010) On this way, it is obtained that this approach may be not conventional in all situations as the influence of alternative feed and also the species of insects on the fatty acid profile.(Ramos-Elorduy 2008)

## Conclusions

To wrap up, due to increasing interests to finding new alternative solvents and food source, *homoptera cicadidae magicicada* can be considered as a new source of edible oil due to acceptable physicochemical characterization where can be contributed in industrial applications and fatty acid profile that introduce it as a nutritious source by covering types of healthy fatty acids as the principal fatty acid was oleic following by palmitic acids and high scale of PUFA and High-MUFA as more than 75% of its fatty acids is consisted of unsaturated fatty acids that can play an important role on healthiness. Furthermore, ethanol-aqueous oil extraction method can be considered as a green and eco-friendly method rather than other traditional methods as hexane extraction by providing acceptable yield of extraction and at the same time a high quality edible oil with lower acidity levels and higher Iodine Value.

## Notes

### Competing Interest Statement

The authors have declared no competing interest.

